# Nanoscale engagement and clusterization of Programmed death ligand 1 (PD-L1) in the membrane lipid rafts of Non-Small Cell Lung Cancer cells

**DOI:** 10.1101/2022.08.09.503318

**Authors:** Martina Ruglioni, Simone Civita, Tiziano Salvadori, Sofia Cristiani, Vittoria Carnicelli, Serena Barachini, Iacopo Petrini, Irene Nepita, Marco Castello, Alberto Diaspro, Paolo Bianchini, Barbara Storti, Ranieri Bizzarri, Stefano Fogli, Romano Danesi

## Abstract

Programmed death ligand 1 (PD-L1) plays a key role in several human tumors, and it has recently become a crucial target for cancer therapy. Yet, its subtle regulatory roles beside the well characterized immunosuppression in PD-1/PD-L1 immune checkpoint are still obscure and should be strictly related to its molecular properties in different cell settings. In this study we targeted the plasma membrane organization of PD-L1 in a Non-Small Cell Lung Cancer (NSCLC) cell line by a multiscale fluorescence imaging toolbox that included super-resolution microscopy approaches to reach the nanoscale. Our findings revealed for the first time that PD-L1 is prevalently engaged in the cholesterol-enriched “raft” regions of the cell membrane. Clusterization and engagement in membrane rafts may afford novel targets for the immuno-oncology strategies in the framework of NSCLC and, possibly, other tumor types.

## INTRODUCTION

Programmed cell death 1 ligand (PD-L1, otherwise known as CD274 and B7-H1) is a key player protein in immune regulation, taking part in one of the most important immune checkpoints of our organism [1]. PD-L1 suppresses T-cell immunity by interacting with specific receptors on T-cells such as PD-1 and CD80. This inhibitory activity unfolds through a complex mechanism that conveys ligand-receptor binding to the silencing of T-cell cytokine production and the induction of anergy and apoptosis [2]. The primary function of immune checkpoints is to maintain the immune homeostasis of the host in physiological conditions [3]. Yet, evasion of the immune system is a recognized hallmark of cancer [4]. Accordingly, several types of cancers exploit the PD-L1/PD-1 interaction to mask cell dysregulation from the T-cell recognition, preventing cancer cell removal [5; 6]. This has led in the last few years to the momentous success of cancer immunotherapy, in which inhibitors (mostly monoclonal antibodies) of the PD-L1/PD-1 binding interaction partially restore the T-cell immune activity and lead to the regression of the neoplasia [7]. For example, most patients with metastatic non-small cell lung cancer (NSCLC) are treated with first-line PD-1 or PD-L1 antibodies, and some of them have prolonged survival [8]. Unfortunately, for several cancers including NSCLC, most patients do not show durable remission and progressively develop resistance to the checkpoint blockade drugs [9]. In this context, the full comprehension of the subtle molecular details underlying the activity of the PD-L1/PD-1 immune checkpoint, particularly those related to the structural properties of the two molecules, appears utterly necessary to provide new rationale for the development of 2nd generation checkpoint blockade strategies. Additionally, it is increasingly clear that PD-L1 has intrinsic regulatory functions to promote the progression of tumors, including the regulation of tumor cell metabolism, sustainment of drug resistance, and the modulation of genome expression upon relocation to the nucleus [10]. Yet, the mechanisms of intrinsic tumorigenic activity of PD-L1 are still largely obscure, urging a better comprehension of the molecular properties of PD-L1 in the cellular settings.

Structurally, PD-L1 is a 40 kDa type-1 transmembrane protein that consists of IgV-like and IgC-like extracellular domains, a hydrophobic transmembrane domain, and a short cytoplasmic tail (cytoplasmic domain, CD) spanning about 30 amino acids [11]. Of particular relevance is the palmitoylation site at residue Cys^272^ of CD [12]. Palmitoylation consists of the covalent attachment of palmitic acid to amino acids (mostly cysteine). This post-translational modification of several membrane proteins is catalyzed by a family of aspartate-histidine-histidine-cysteine (DHHC) acyltransferases. It has been demonstrated that PD-L1 palmitoylation by DHHC3 in colon cancer stabilizes PD-L1 by suppressing CD ubiquitination, which in turn triggers lysosomal degradation of the immune checkpoint ligand [13]. Analogous stabilization of PD-L1 by zDHHC9-mediated palmitoylation was revealed in breast cancer [14]. Accordingly, specific mutation of Cys^272^ or inhibition of the palmitoyltransferase was able to restore immune T-cell activity against cancer cells [14]. These findings led to the palmitoylation of PD-L1 (and more recently of PD-1) being a new target for immuno-oncology [15; 16]. Palmitoylation of membrane proteins has a widespread role in cell biology to increase their affinity to the lipid raft regions of the plasma membrane (PM) [17–19]. A simplified-yet scientifically fertile-model of PM invokes an interlaced combination of liquid-order (L_O_, also referred to as ‘‘lipid raft’’) and liquid-disorder (L_d_) nanophases, enriched respectively in saturated and unsaturated lipids, together with different amounts of cholesterol [20–22]. The extension of these nanophases is modulated by the confining action of the cytoskeleton and it is utterly dynamic, as it leverages a continuous exchange of proteins and protein complexes [23]. This paradigm of membrane assembly was proposed to be a crossroad in every membrane process, such as the formation of protein clusters, signal transduction, endocytosis, and cell polarization and motility [23]. Of note, the structure of the raft can be planar (rectilinear) or non-planar (invaginated). Non-planar rafts can be largely identified with caveolar regions of the PM, flask-like invaginations of the plasma membrane enriched in the protein Caveolin-1 that contribute to several aspects of cell physiology, including signal transduction events and the specialized caveolar endocytosis mechanism [24; 25].

In spite of the palmitoylation at CD, it is unclear whether PD-L1 is really engaged in lipid rafts, albeit an increased raft localization of this protein seems to take place in melanoma cell lines stimulated with an anti HLA-DR antibody [26]. Additionally, a recent preprint carried out *in vitro* experiments on giant plasma membrane vesicles (GPMV) to determine the partition of PD-L1 into the lipid-ordered microphase [27]. In this study, the authors found that palmitoylated PD-L1 displays significant affinity to membrane-ordered phase. Yet, the loss of cytoskeletal organization in GPMV may prevent a correct assessment of the PD-L1-raft relationship in physiological cell conditions. Beside putative raft engagement, it is worth noting that no notion on the actual membrane organization of PD-L1 is available.

In this work, we set out to investigate the PD-L1 organization on the PM of epithelial growth factor receptor (EGFR) mutant NSCLC cells by a multiscale toolbox of fluorescence microscopy techniques. More specifically, we first leveraged sensitive imaging systems such as Total Internal Reflection Fluorescence (TIRF) and Image Scanning Microscopy (ISM) to reveal the partition of PD-L1 in raft regions by colocalization measurements with molecular hallmarks of lipid rafts. Then, we characterized the nanoscale morphology of PD-L1 on the PM by using single-molecule localization microscopy (SMLM) according to the dSTORM imaging scheme [28]. Both ISM and dSTORM belong to the family of super-resolution imaging techniques that recently revolutionized the fluorescent microscopy scenario [29]. ISM reaches the 100-140 nm resolution just by a smart arrangement of the detector units [30]. dSTORM leverages the ability to randomly activate the fluorescent emission of only a small subset of fluorophores, in order to distinguish them spatially and enabling spatial resolution as low as a few nanometers [28]. By this approach, we demonstrated that PD-L1 organizes into nanoclusters on the PM likely upon engagement with membrane rafts.

## MATERIALS AND METHODS

### Reagents, plasmids and antibodies

Chemical and biological reagents were purchased from Sigma-Aldrich/Merck if not-otherwise specified.

The PD-L1-EGFP plasmid (pEGFP-N1/PD-L1, [31]) was a gift from Prof. Mien-Chie Hung of The University of Texas - M. D. Anderson Cancer Center through the Addgene plasmid repository (Addgene plasmid #121478). Caveolin-1-EGFP (Cav1-EGFP) was a kind gift of Prof. Ari Helenius of ETH Zurich and it has been described in [32]. Glycosyl phosphatidylinositol-EGFP (GPI-EGFP) was a kind gift of Prof. Jennifer Lippincott-Schwartz of Howard Hughes Medical Institute’s Janelia Research Campus and it has been described in [33].

The following antibodies were used throughout this work:

- αPD-L1-647: Rabbit anti-PD-L1 monoclonal IgG conjugated to AlexaFluor647 ( D8T4X, #41726, Cell Signaling, Euroclone, Milan, Italy). Immunolabeling dilution: 1/100 in PBB
- αPD-L1: Rabbit anti-PD-L1 monoclonal IgG (mAb D8T4X, #86744 Cell Signaling, Euroclone, Milan, Italy). Immunolabeling dilution: 1/400.
- αACE2-647: Rabbit anti-ACE2 monoclonal IgG conjugated to AlexaFluor647 (EPR24705-45, #283658 AbCam, Prodotti Gianni, Milan, Italy). Immunolabeling dilution: 1/50.
- αr647: donkey anti-rabbit monoclonal IgG conjugated to AlexaFluor647 (a31573, ThermoFisher, Milan, Italy). Immunolabeling dilution: 1/400.
- αr647: donkey anti-rabbit monoclonal IgG conjugated to AlexaFluor488 (a21206, ThermoFisher, Milan, Italy). Immunolabeling dilution: 1/400.
- i647: rabbit IgG Isotype Control AlexaFluor647-conjugated (DA1E, #2985, Cell Signaling Technologies). Immunolabeling dilution: 1/100.

### Cell culture

EGFR-mutated NSCLC cells (H1975, H3255, H1650, PC9, and HCC827) cells were cultured at 37°C in the presence of 5% CO_2_ in RPMI-1640 medium containing phenol red and supplemented with NaHCO_3_, 2 mM L-glutamine, 1% of sodium pyruvate, 1% of penstreptomycin, and 10% of fetal bovine serum (FBS).

African green monkey kidney VeroE6 cells were cultured at 37 °C in the presence of 5% CO_2_ in DMEM high glucose medium supplemented with heat-inactivated 10% FBS, 2 mM L-glutamine, 10 U/ml penicillin, and 10 mg/ml streptomycin.

Cells were seeded (3×10^5^) in 35 mm glass-bottom dishes (WillCo Wells BV, Amsterdam, The Netherlands) with 2 ml of culture medium and maintained at 37°C and 5% CO_2_ for 24 prior to transfection (*vide infra*) or immunostaining.

### Cholesterol depletion

Cell viability upon methyl-beta-cyclodextrin (MBCD) exposure was evaluated by a MTT (3-(4,5-dimethylthiazol-2-yl)-2,5-diphenyltetrazolium bromide) test in a 96-well plate according to the manufacturer’s (Merck) protocol. Absorbance was read by using a Plate Reader (BIOTEK FLx800, Agilent Biotek, Cernusco sul Naviglio, Milan, Italy). Cell viability was expressed through the viability ratio (vr=100*sample absorbance/control absorbance) as 100-vr. Cell incubation with 5 mM MBCD at 37@ and 5% CO_2_ for 1h gave cell viability > 50% and this MBCD concentration was thereafter applied to cells.

Transient expression of Caveolin-1-EGFP, GPI-EGFP and PD-L1-EGFP in HCC827 cells

A solution made up of 1 µL transfection plasmid (Caveolin-1-EGFP, GPI-EGFP, or PD-L1-EGFP, concentration: 1 µg/µL) in 100 µL Opti-Mem reduced Serum Medium (Thermofisher, Milan, Italy) was added to a solution of 3 µL Lipofectamine 2000 (Thermofisher, Milan, Italy) in 100 µL Opti-Mem. The resulting mix was incubated for 5 minutes at room temperature and then added to HCC827 cells adhered to 35 mm glass bottom Willco (WillCo Wells BV, Amsterdam, The Netherlands) dishes. Cells were then incubated at 37@ with 5% CO_2_ for 24-38 hours prior to immunostaining.

### Immunostaining of cells

Adherent cells (untransfected and transfected) in WillCo dishes were fixed with paraformaldheyde (PFA) 2% and glutaraldehyde (GLU) 0.2% in PBS (pH 7.2) for 15 min at RT, rinsed three times with PBS. Cells were then exposed to glycine 0.1 M in PBS for 30 min and rinsed 3 times with PBS and 3 more times with 0.5% Bovine Serum Albumin (BSA) in PBS (PBB). Subsequently, cells were maintained for 40 min at RT in 2% BSA in PBS and rinsed three more times with PBB.

For direct immunostaining, HCC827 cells were incubated for 1h with αPD-L1-647 or i647 diluted in PBB. Transfected VeroE6 cells were incubated for 1h with αACE2-647 in PBB.

For indirect immunostaining, untransfected HCC827 cells were incubated for 1h with αPD-L1 diluted in PBB. After rinsing three times with PBB, cells were incubated for one more hour with αr647 diluted in PBB. Cells were then extensively rinsed with PBS.

As positive control for colocalization experiments, cells immunostained by αPD-L1-647 were counterstained for 1h with αr488 diluted in PBB and then extensively rinsed with PBS.

When nuclear staining was needed, immunostained cells were exposed for 5 min to 1 mg/100 ml Hoechst 33342 (Thermo Fisher Scientific, Waltham, Massachusetts) in water.

### Determination of the labeling degree of anti-PD-L1 conjugated to Alexa647

The labeling degree of αPD-L1-647 was determined by the classical procedure developed by Molecular Probes^1^. In more details, the absorbance of a solution of the antibody in PBS (pH 7.2) was collected in the 250-800 nm wavelength range by means of a Jasco V-550 UV-Vis Spectrophotometer (Jasco, Milan, Italy). Molar concentration of the antibody was calculated by the following equation:

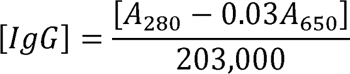

where *A*_280_ and *A*_6s0_ are the absorbances at 280 and 650 nm, respectively. The concentration of AlexaFluor647 fluorophore was calculated by:

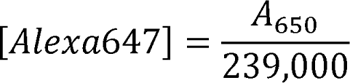

Finally, the labeling degree D was calculated simply as:

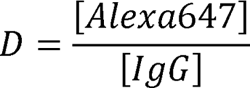

We found out: D = 1.09.

### Flow cytometry

Flow cytometry was carried out by a MACSQuant Flow Cytometer (Miltenyi Biotec instruments, Bergisch Gladbach, Germany). Viable cells (5×10^5^) were resuspended in 100 µl MACSQuant running buffer. Cells were then labeled with either αPD-L1-647 (sample) or i647 (Isotype Control), at the proper dilution (*vide supra*) for 15 min at 4°C. Unlabeled cells were used as autofluorescence (blank) control. After washing to remove the excess unbound antibody, cells were resuspended in 450 µl of the running buffer and filtered to eliminate any aggregates.

The flow cytometer was set using cells stained with isotype controls. Frequencies of cell populations were calculated on total events, after exclusion of cell debris on FSC (forward scatter) vs. SSC (side scatter) density plots and doublets on FSC-A vs. FSC-H. Quantification of PD-L1 expression in tested cell lines was carried out by comparing the signals of sample, isotype control, and blank cells. Acquisition was performed collecting 10,000 events that were analyzed by MACSQuant® Flow Cytometer using the MACSQuantify® Software (Miltenyi Biotech, Bologna, Italy). Statistical significance was assessed by the Anova Test with Bonferroni correction.

### Confocal microscopy

Fluorescence was measured by a laser scanning confocal microscope Nikon, Eclipse Ti. Samples were viewed with a 63x Apochromat NA=1.4 oil-immersion objective. Pixel dwell time was adjusted to 1.52 µs and either 512×512 pixel or 1024×1024 images (line scan speed was changed accordingly) were collected (line average factor: 4). The acquisition channels were set as follows:

- blue (Hoechst 33342): λ_ex_=405 λ_em_= 450/50 nm;
- green (AlexaFluor488/EGFP): λ_ex_=488, λ_em_= 525/50 nm;
- far-red (AlexaFluor647): λ_ex_=640, λ_em_= Long Pass 650 nm.

The pinhole size was set to 1 airy unit (AU) for the green acquisition channel (0.87 AU for the Far-red channel). Images were visualized and processed by the open-source software Fiji (NIH, Bethesda).

### Total internal reflection microscopy

Imaging in Total Internal Reflection mode (TIRF-M) was carried out by a Leica AF6000 fluorescence microscope equipped with a TIRF-M condenser and 100x oil-immersion objective (NA 1.47). We adjusted the penetration depth of the evanescent wave to 100-150 nm. Fluorescence was recorded by a cooled EM-CCD (Hamamatsu C1900–13). The microscope was equipped with laser lines for excitation of Alexa488/EGFP (488 nm) and AlexaFluor647 (640 nm). The acquisition channels were set as follows:

- green (AlexaFluor/EGFP): excitation was set to 488 nm, the emission was collected between 520 and 550 nm, and a dichroic filter at 502 nm separated excitation from emission;
- far-red channel (AlexaFluor647): excitation was set at 640 nm, emission was collected by a superposition of 600/40 641/75 and LP650 filters (yielding light at 604–620 and 650–679 nm), and a dichroic filter at 570 nm separated excitation from emission.

### Image Scanning Microscopy

Images in the green (AlexaFluor488/EGFP) and far-red (AlexaFluor647) channels were acquired by a custom-made setup as described in [34]. Pixel reassignment and Lucy-Richardson image deconvolution (ISM^++^) were implemented exactly as described in [30].

### Colocalization analysis

Functional colocalization of images collected in the green (AlexaFluor488/EGFP) and far-red (Alexa647) channels was quantified by Pearson’s coefficient R using the JACoP plugins of Fiji [35], which implements the automatic thresholding approach developed by Costes et al. [36].

### Super-resolution microscope setup and imaging

An N-STORM Eclipse Ti2 microscope (Nikon Instruments) equipped with an oil immersion objective (CFI Apo TIRF 100×, NA 1.49, oil; Nikon) was used to acquire 20,000-80,000 frames with 30 ms acquisition time using TIRF illumination. Excitation intensities after the objective were as follows: 35.4 mW for the 647 nm read-out (450 mW laser; Nikon Instruments) and 5.5 mW for the 405 activation (450 mW laser; Nikon Instruments). We set a repeating cycle of 1 activation frame at 405 nm / 1 readout frames at 647 nm. Image detection was performed with a cMOS camera (Orca-Flash 4.0 C13440, Hamamatsu). The Perfect Focus System (Nikon) was used during the entire recording process. The fluorescence-emitted signal was spectrally selected by the four-color dichroic mirrors (ZET405/488/561/647; Chroma) and filtered by a 4-bandpass filter (ZT405/488/561/647; Chroma). For imaging, cells were embedded in the Everspark™ buffer (Idylle, Paris, France).

### Single-molecule localization and filtering

The stacks were processed by Thunderstorm, a Fiji plugin for PALM and STORM data analysis. In the “Camera setup” menu, we set: pixel size = 162 nm, photoelectrons per A/D count: 0.49, base level=100 A/D count. The localization algorithm (“Run analysis”), was carried out by setting the following parameters: a) pre-filter: wavelet filter (B-spline), scale: 2, order: 3; b) approximate localization of molecules by “local maximum” method with 3x standard deviation of the 1st wavelet level as threshold and 8-neighborhood connectivity; c) sub-pixel localization by the Integrated Gaussian method, performing maximum likelihood fit with initial sigma 1.6 pixels and fitting radius 3 pixels.

Next, we cleaned the obtained results from drift and localizations not lying on the focal plane by the following post-filtering algorithm: a) drift correction by correlation (3-5 points); b) removal of localizations with sigma >180 nm OR uncertainty > 20 nm OR intensity>5000 photons. The average of the localization precision means over all datasets was found to be < xy >=9±1 nm (average±SE).

Finally, multiple blinking signals from the same fluorophore were condensed into a single localization by the Distance Distribution Correction (DDC) algorithm [37]. In short, DDC relies on the pairwise distribution of the distance of localizations Z with a lag time much longer than the average fluorescence survival time, and it applies to any kind of blinking fluorophore. DDC only needs as input two parameters, obtainable by two subroutines of the code: 1) the width of the bins for the histograms of the probability distributions (called “bin resolution”, approximately 2x the average localization precision of the localization dataset) and 2) the time after which the blinking fluorophore is bleached off (survival time). For our data, DDC led to a “bin resolution” 20 nm. The survival time of AlexaFluor647 was set to 100 frames (3 s) as the Z vs. frame curve leveled off at these values.

Graphical renderings of the processed single localization data (dSTORM images) were generated by using gaussian functions peaked on each localization and with sigma corresponding to the relevant localization precision and setting the pixel size to 10-30 nm.

### Ripley’s and G&F analysis

A custom-made MATLAB (version R2022a, Mathworks, Natvick, MA, USA) code was developed to compute Ripley’s H function in an unlimited a priori rectangular area. This code first calculates Ripley’s K(r) function according to:

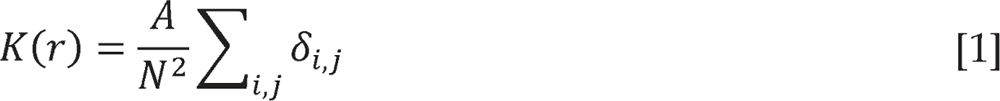

where A is the area of the selected rectangular ROI in the dSTORM image, *N* is the number of total localizations in the area, r is the spatial variable (radius) that determines the *K* function, and O_i,j_ = 1 if the distance between the *i*-th and *j*-th localizations is less than *r*, otherwise O_i,j_ = 0.

*K*(r) is then transformed into the function L(r) by:

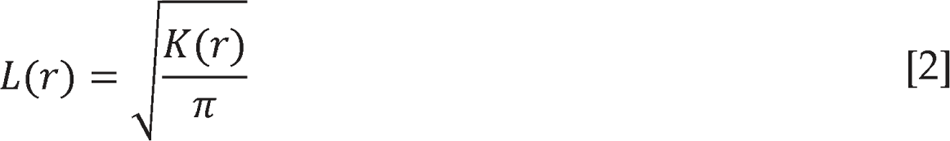

For a perfectly isotropic distribution, *L(r) = r*. Thus, we can define the Ripley’s function *H(r) = L(r) - r* to measure the spatial randomness of the localization map. According to its definition, *H(r)* must be zero when the localizations are distributed isotropically, and positive when particles are clustering. The *r_m_* value at which *H(r)* According to its definition, H(r) must be zero when the localizations are distributed reaches a maximum typically lies between the average cluster radius (*r_c_*) and the average cluster diameter (2 *r_c_*), depending on the mean distance between the clusters within a factor of 1.3 from *r_m_* [39]. Our code computed *K(r), L(r)*, and *H(r)* for 0 < r ≤ 1 [38]. Yet, for a random distribution of clusters, the cluster radius can be estimated µm, automatically identifying the value *r = r_m_* corresponding to the maximum of *H(r)*. Of note, the code used toroidal edge correction to comply with the ROI boundaries [40]. To estimate the effect of precision localization on the Ripley’s function H(r), we adopted displacements *e_X_* and *e_Y_* to the X and Y coordinates of each position, respectively. *e_X_* a Monte Carlo approach. For each dataset we generated 25 artificial datasets adding two and *e_Y_* were randomly taken from a Gaussian distribution with average zero and standard deviation equal to <XY>/√2, where the square root considers the splitting of deviation of the calculated 25 *r*_m_ values was considered as the *_erm_* error of the the radial precision into the two orthogonal coordinates X and Y. The standard unperturbed original dataset.

The Ripley analysis was used to build a spatial binary map of the blinking molecules belonging to clusters according to the procedure developed by Getis and Franklin (G&F specifically, we set a *r* = 40 nm (the closest integer to the average *r_c_*, see Results section), analysis, [41]) and recently applied by Kennedy et al. to STORM data [42]. More and all molecules characterized by *L*(*r* = 40 nm) > 90 nm (i.e. more than two-fold the value pertaining to isotropic distribution at that radius) were replaced by an intensity-uniform disk. Such a disk was centered in the localization coordinates and had a radius *r* = 30.8 nm, corresponding to the minimum cluster radius under the assumption of the maximum conversion factor 1.3x between the spatial metrics of Ripley function and the the average number of localizations per cluster (d_c_), as well as the % of overall actual size of a cluster (*vide supra*). The morphological analyses of binary maps afforded localizations (*f*) that are found in clusters.

### Graphics and statistics

Graphs were prepared using Prism 7 (GraphPad Software, Boston, MA, USA) and IgorPro8 (Wavemetrics, Lake Oswego, OR, USA) software. Histograms are shown as the mean +/- standard deviations. Statistical analysis was performed by Prism 7.

## RESULTS

### Selection of the NSCLC cell line model

The preferred initial treatment of EGFR-mutant NSCLC is the use of EGFR tyrosine kinase (TKI) inhibitors, as these patients exhibit poor response to anti-PD-1/PD-L1 treatments. However, acquired drug resistance may limit the long-term efficacy of EGFR TKIs and no further effective treatment options are currently available for these patients [43]. From literature data, it is known that activation of EGFR is positively associated with the activation of PD-L1 in NSCLC cells [44]. Accordingly, we set out to inspect the expression level of a family of NSCLC cell lines with different mutation patterns of the EGFR gene, namely H1975, H3255, H1650, PC9, and HCC827 [45]. All these cell lines are characterized by a noticeable transcription level of the PD-L1 gene as assessed from the cancer cell line encyclopedia (Table 1).

**Table 1:** Mutation pattern and transcription level of several EGFR-mutated NSCLC cell lines from the cancer cell line encyclopedia^2^

|  | H1975 | H3255 | H1650 | HCC827 | PC9 |
| --- | --- | --- | --- | --- | --- |
| EGFR mutation | ex20 T790M +<br>ex21 L858R | ex21 L858R | ex19<br>E746_A750del | ex19<br>E746_A750del | ex19<br>E746_A750del |
| mRNA(rpkm) | $5.41 \cdot 10^5$ | $7.03 \cdot 10^5$ | $1.74 \cdot 10^5$ | $1.74 \cdot 10^5$ | $2.09 \cdot 10^5$ |

The actual expression level in the cells was evaluated by flow cytometry taking advantage of a fluorescently labeled rabbit IgG that selectively targets an extracellular epitope of PD-L1 (αPD-L1-647) extracellularly, thus enabling direct immunolabeling of non-permeabilized cells with full membrane selectivity. The use of a rabbit AlexaFluor647-labeled IgG isotype (i647) provided the necessary negative control. Analysis of the histograms obtained from flow cytometry revealed the positive immunostaining of PD-L1 as compared to the negative control (**Figure 1a**). Comparison of the median values of cytofluorograms hinted to the lowest expression of membrane PD-L1 in H3255, albeit the observed fluorescence value was still significantly higher than the negative control (**Figure 1b**). HCC827 resulted the cell line with highest PD-L1 expression, and it was selected as the study cell model to enable the largest sensitivity in the subsequent imaging experiments.

**Fig. 1.**
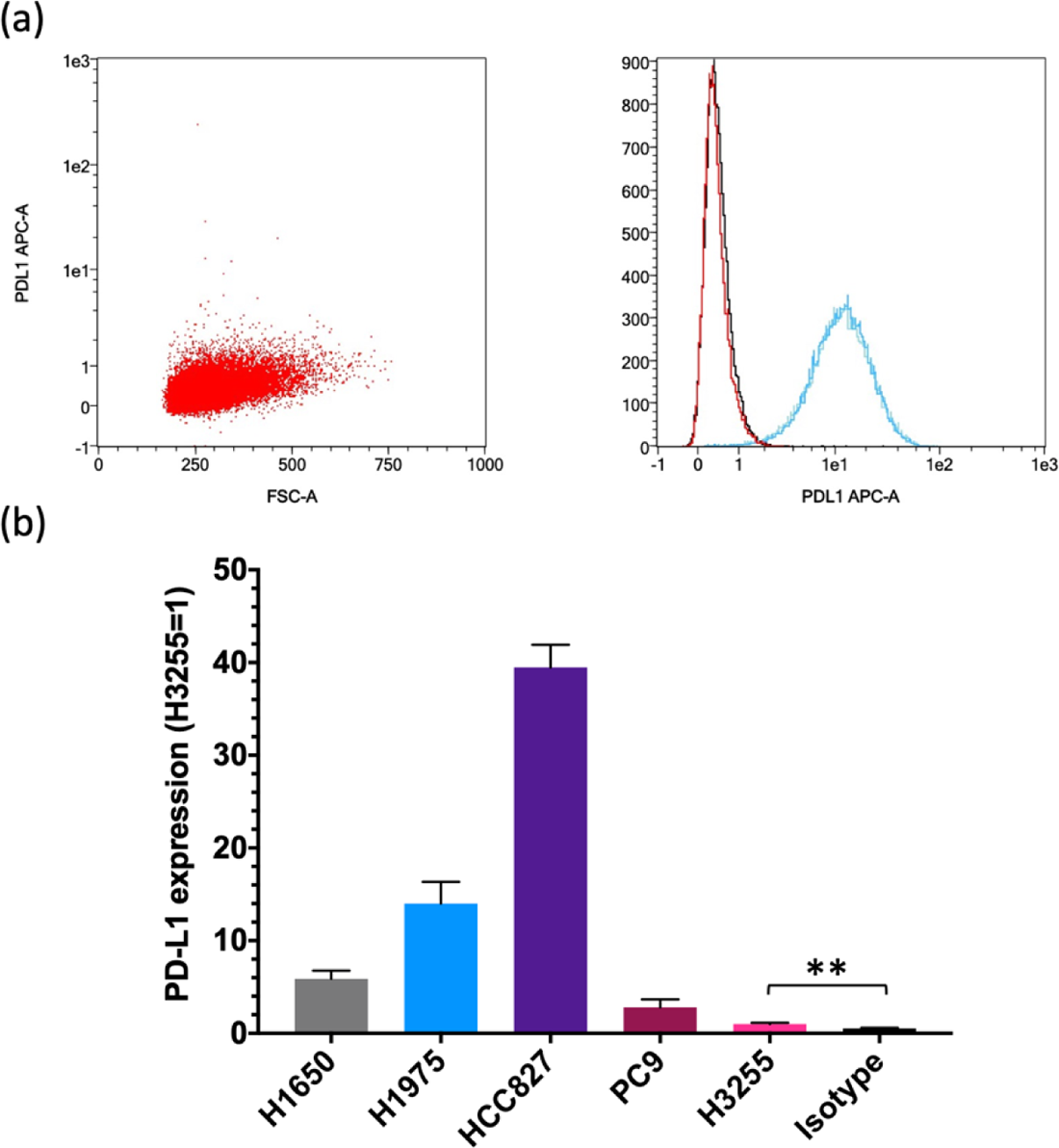
Expression of PD-L1 in NSCLC cell line models bearing mutated EGFR. (a) Representative flow cytometry dot plot and frequency histograms showing immunostaining of PD-L1 in HCC827; non labeled cells (black), IgG Isotype Control (red), positively immunostained cells (blue). (b) Relative PD-L1 expression (PD-L1**r**) in five cell lines (we assumed PD-L1r = 1 for H3255) H1650: PD-L1r = 5.9±0.9 (average±SD); H1975: PD-L1r = 14±2; HCC827: PD-L1r = 39±2; PC9: PD-L1r = 2.8±0.9; Isotype: PD-L1r= 0.5±0.1. Student’s t-test: p<0.05 (**). Imaging of membrane PD-L1 and cholesterol depletion αPD-L1-647, or its non-labeled version αPD-L1, were used to stain PD-L1 on the outer surface of PM by direct or indirect immunofluorescence, respectively. Confocal and TIRF microscopy images highlighted the heterogeneous distribution of PD-L1 in submicron foci on the PM, allegedly positing the existence of aggregate structures (**Figure 2a,b**). Of note, optical 3D reconstruction of several focal planes acquired by confocal microscopy confirmed the almost exclusive PM staining of PD-L1 (**Figure 3**).

**Fig. 2.**
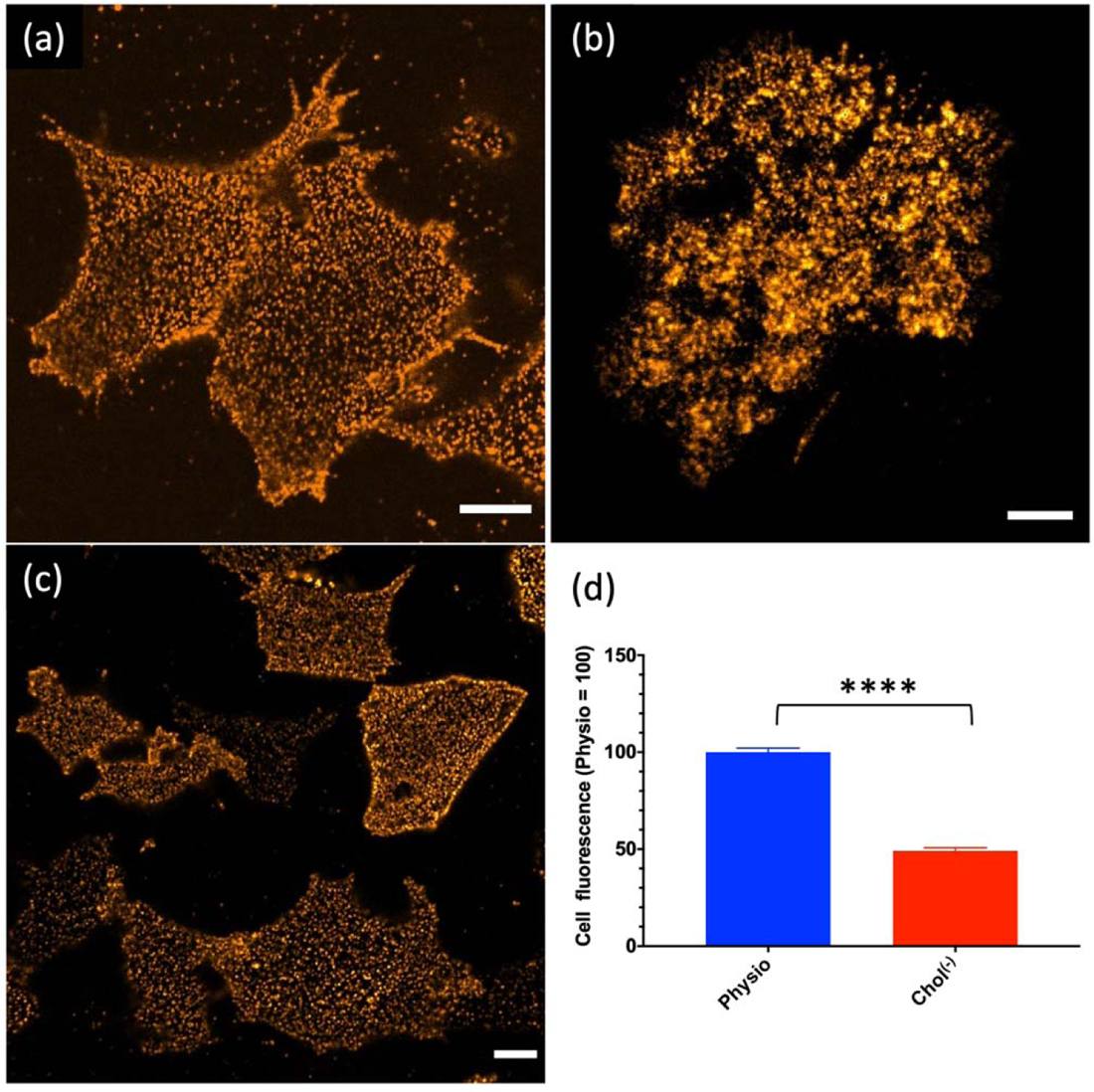
Confocal and Total Internal Reflection Fluorescence (TIRF) imaging of PD-L1 in HCC827 cells upon indirect immunostaining of PD-L1 and using AlexaFluor647 as fluorescent reporter. (a) Confocal image of HCC827 cells in physiological conditions (Physio cells). (b) TIRF image of Physio cells. (c) Confocal image of HCC827 upon cholesterol depletion by 5 mM MBCD for 1h at 37 °C (Chol^(-)^ cells). (d) Comparison between the fluorescence of Physio and Chol^(-)^ cells (Physio = 100); p<0.0001 (****). Scale bar: 10 μm. Cholesterol depletion by methylbetacyclodextrin (MBCD) is a classical way to disrupt membrane rafts [46] and thereby modulate the residence of raft-engaged proteins [47]. Accordingly, we exposed HCC827 cells to the minimum amount of MBCD to preserve >50% viability, as judged by the MTT assay (Chol^(-)^ cells: 61% viability). Confocal images of Chol^(-)^ cells revealed that cholesterol depletion did not affect the membrane organization of PD-L1 into submicron foci (**Figure 2c**). Yet, a statistically significant fluorescence reduction (w50%) upon cholesterol depletion was observed in comparison to non-depleted cells (**Figure 2d**, p<0.0001).

**Fig. 3.**
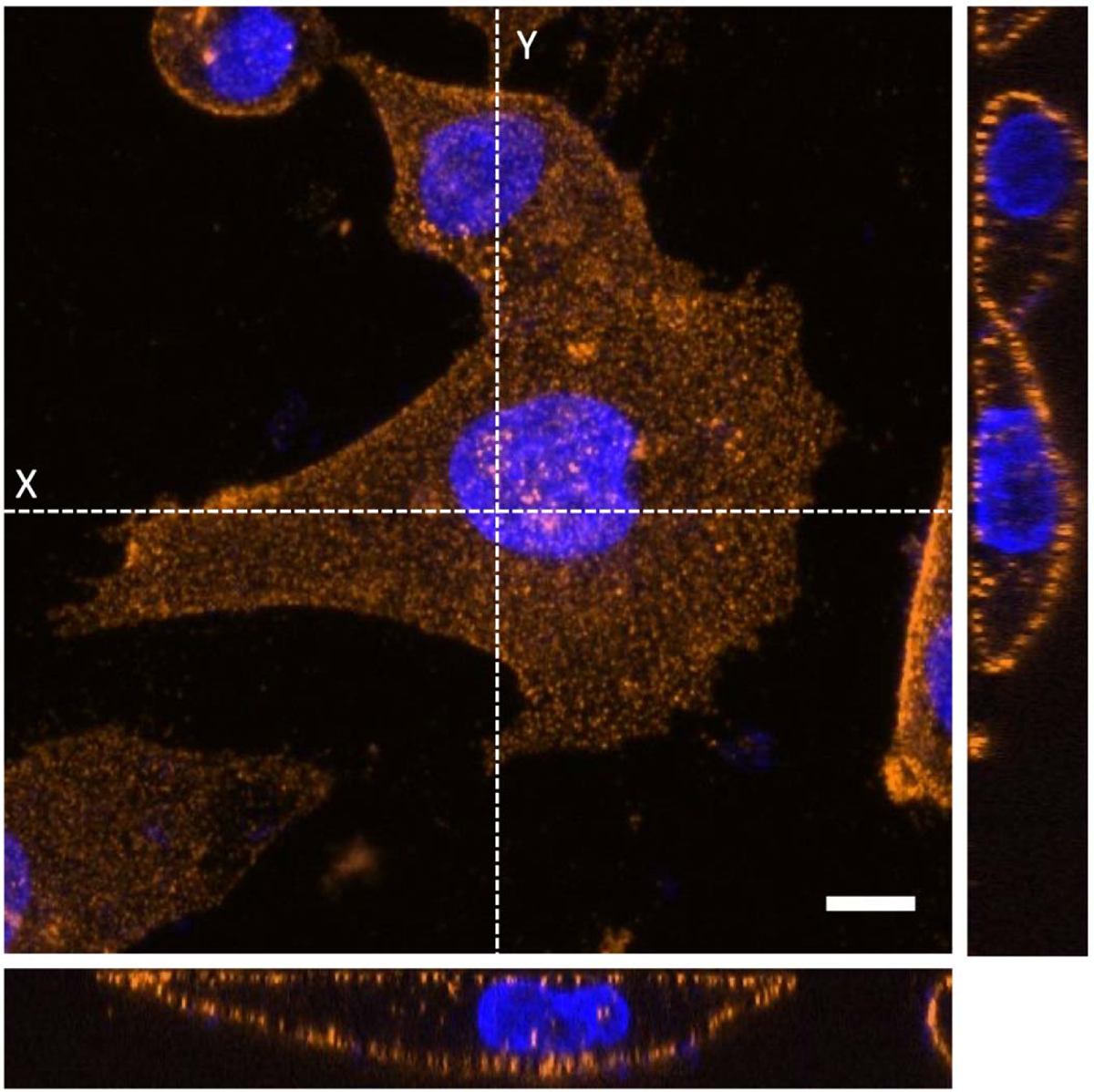
Spatial distribution of PD-L1 on HCC827 cells by confocal microscopy. Central image is a maximum projection of a stack of images collected over 13.4 μm along the Z-direction. Left and bottom panels show the YZ and XZ images collected by sectioning along the Y and X dashed lines, respectively. Blue: Hoechst, orange: PDL1. Scale bar: 10 μm.

### Imaging of membrane PD-L1 and colocalization with lipid raft markers

To investigate the relationship between PD-L1 and lipid rafts, we carried out colocalization studies of directly immunostained PD-L1 with two EGFP-labeled proteins acting as raft hallmarks, namely glycosylphosphatidylinositol (GPI) or Caveolin-1 (Cav-1). GPI is a common anchoring motif of several proteins to the cell membrane and GPI-anchored proteins were shown to reside mostly in lipid rafts [17; 48]. Cav-1 is the hallmark of caveolae, flask-like invaginations of the plasma membrane which constitute a subset of lipid membrane rafts and are responsible for “caveolar” endocytosis [24; 49]. As positive colocalization control (CTRL^+^), we used an anti-rabbit IgG fused to AlexaFluor488 (αr488) to counterstain the primary label αPD-L1-647. Of note, AlexaFluor647 is characterized by almost no emission overlap with EGFP, thereby avoiding spurious colocalization effects due to spectral overlap.

Visible yellow patches in the merged TIRF images showed extended colocalization between the raft markers (colored in green) and PD-L1 (colored in red) (**Figure 4a-f**). This pattern was quantitatively confirmed by the calculation of Pearson’s coefficient, R, which measures the stoichiometric correlation between the two fluorescent partners as a proxy of their functional association [50]. The collected R values (**Figure 5**) showed that PD-L1 functionally colocalizes with both GPI-EGFP (<R>=0.37±0.04, average±SE) and, to a slightly lesser extent, with Cav-1 (<R>=0.31±0.05). In both cases, colocalization was not complete, as judged by comparison with CTRL^+^ (<R>=0.59±0.04).

**Fig. 4.**
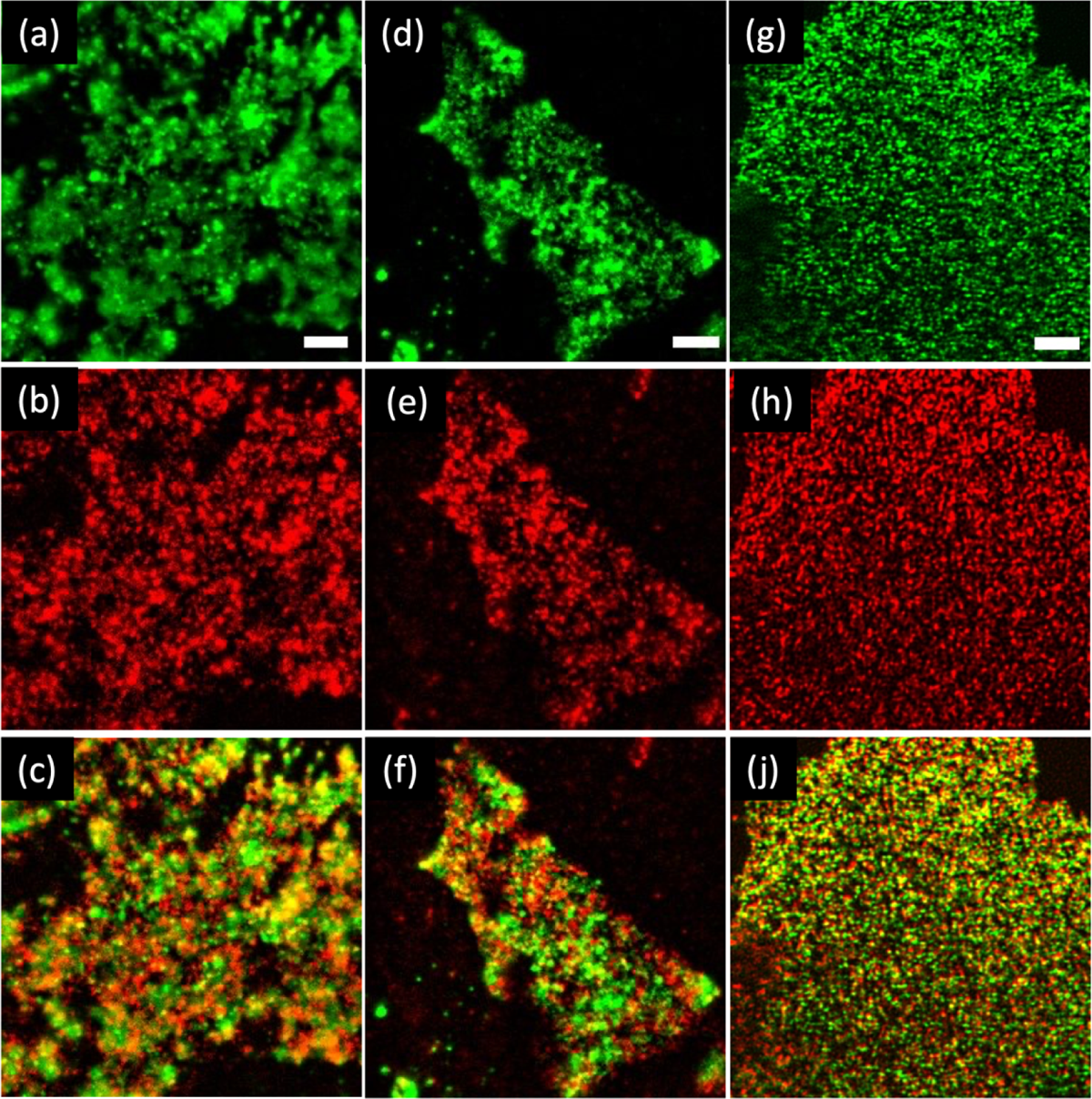
Colocalization of PDL1 with raft membrane markers in HCC827 cells. (a-c) TIRF image of transiently expressed GPI-EGFP (a), immunostained PD-L1 (b), and color merge (c). (d-f) Same as for (a-c), but GPI-EGFP was replaced by Cav-1-EGFP. (g-j) Same as for (a-c), but images were acquired by Image Scanning Microscopy followed by Lucy-Richardson deconvolution (ISM++). Scale bar: 5 μm.

**Fig. 5.**
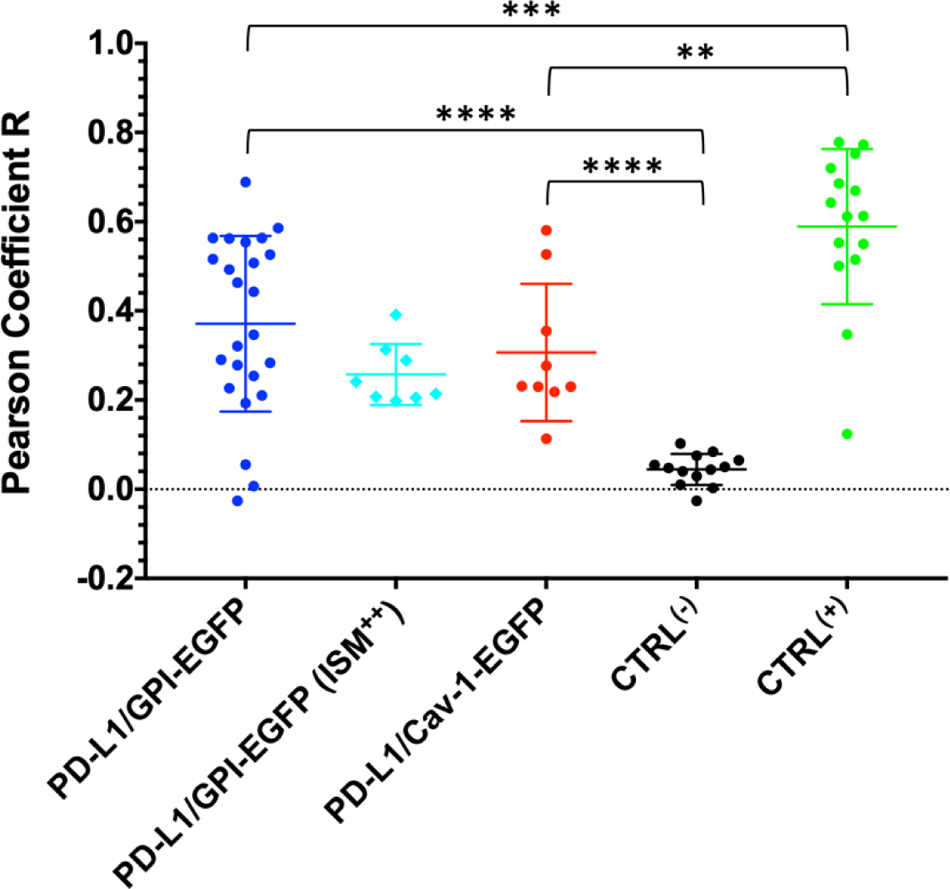
Quantification of PD-L1 colocalization with raft markers by the Pearson’s coefficient R. Circles refer to single measurements while horizontal bars show average±SD. The Mann-Whitney U test was used to compare differences among datasets. p<0.01 (*), p<0.001 (***), p<0.0001 (****).

To provide a meaningful negative control (CTRL^-^), we determined the colocalization of the membrane protein ACE2 with PD-L1 in VeroE6 cells transiently expressing PD-L1-EGFP. Indeed, we recently demonstrated that ACE2 localizes in the non-raft regions of the VeroE6 cells. Remarkably, we found almost no colocalization (<R>=0.04±0.01) fo CTRL^-^, with high statistical significance compared to GPI-EGFP/PD-L1 or Cav1-EGFP/PD-L1 (p<0.0001, **Figure 5**).

At the typical optical resolution of diffraction limited systems such as TIRF or confocal microscope (200-300 nm), the extent of colocalization of nanoscale objects might be partly overestimated [51]. To further verify the consistency of the PD-L1 colocalization in rafts, we carried out Image Scanning Microscopy (ISM) followed by Lucy-Richardson deconvolution (ISM^++^ imaging mode) of the couple GPI-EGFP/ PD-L, adopting an approach we recently leveraged to reveal the colocalization of nanoclustered proteins at 100-160 nm resolution [34]. Note that at such spatial scale we may expect the fluorescence still coming from an ensemble of nanoscale emitters, thus preserving the concept of stoichiometric correlation expressed by the Pearson coefficient. Consistently with the TIRF measurements, ISM^++^ images revealed significant GPI-EGFP/PD-L1 colocalization (**Figure 4g-j**). Quantitatively, we found <R>=0.26±0.02, this being a value almost indistinguishable from the one collected by TIRF (p=0.07, **Figure 5**).

### Nanoscale cluster organization of PD-L1 on cell membrane

The nanoscale arrangement of PD-L1 on the basal membrane of HCC827 cells was investigated by dSTORM in TIRF mode using direct immunolabeling by αPD-L1-647 (**Figure 6**). Operatively, we collected 20-40,000 frames (600-1200 s) by alternating illumination at 640 nm (fluorescence excitation and switching-off) and 405 nm (fluorescence switching-on). Raw localization coordinates obtained by ML method were each dataset showed a localization precision (σ_xy_) distribution ranging between 1 and at first cleaned to compensate for drift and to remove off-focus emitters. After cleaning, all the σ_xy_ distributions, we found <σ_xy_>= 9±1 nm (average±SE, #29 cells). The average 20 nm with mean around 8-10 nm (**Figure 6a, inset**). Taking the average of the means of localization density was 46±16 localization/µm^2^ (average±SE, #29 cells). For comparison, cells treated with the isotype control antibody (i647) were characterized by a much lower localization density (2.3±0.7 localization/µm^2^, average±SE, #8 cells). Thus, we concluded that in our datasets, non-specific labeling was no more than 5%.

**Fig. 6.**
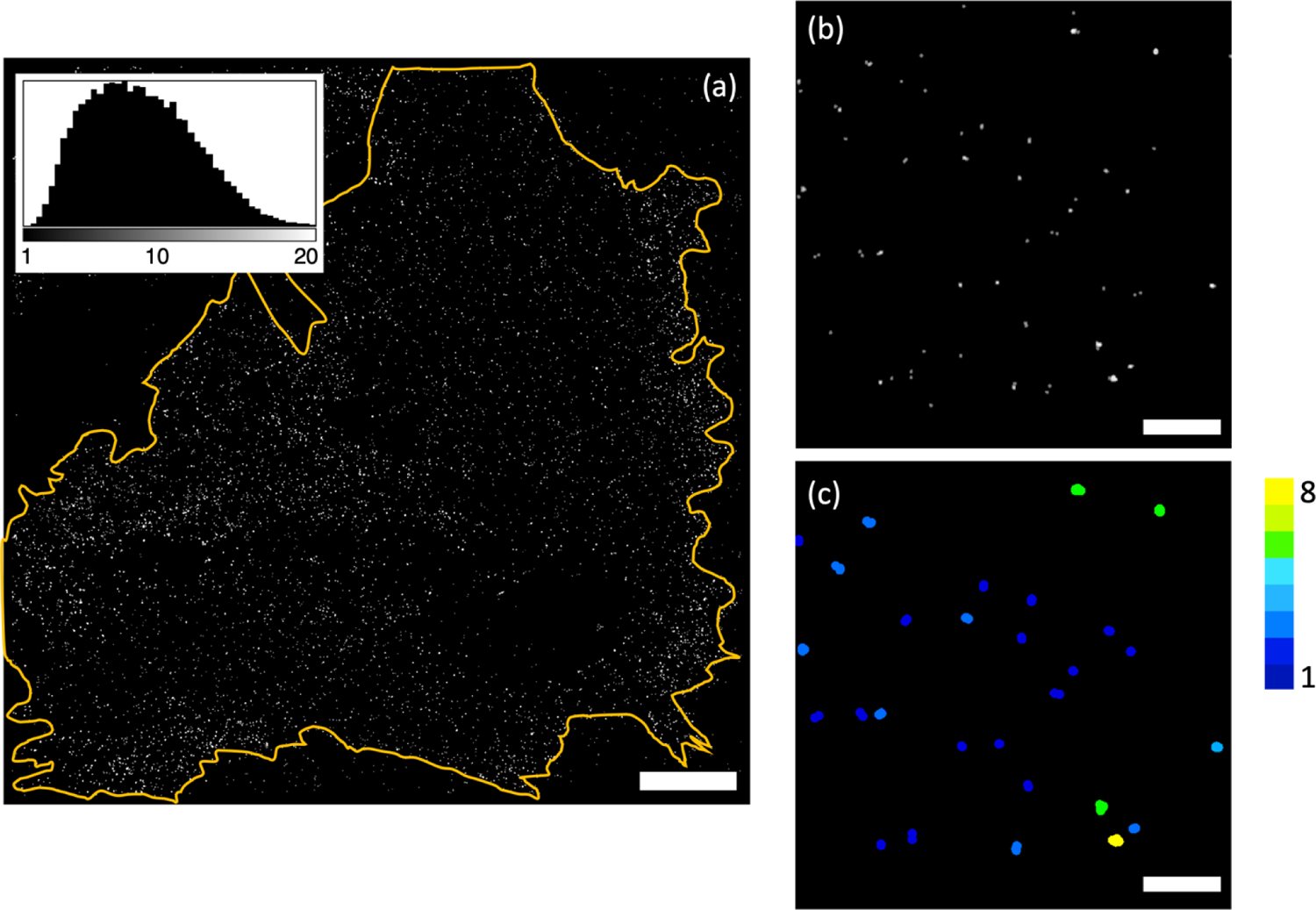
Single molecule localization microscopy (TIRF-dSTORM mode) of PD-L1 on the basal membrane of a HCC827 cell. (a) Gaussian rendering of dSTORM map of one HCC827 cell membrane immunostained for PD-L1. In top left inset is reported the distribution of the localization precisions before DDC correction. (b) Zoom of a 2.8 μm x2.8 μm region of the same cell. (c) Quantitative clustering map of the region enclosed in (b) according to Getis and Franklin’s local point pattern analysis. The pseudo-color scale refers to the number of emitter enclosed in each cluster. Scale bar: 2 μm (a) and 500nm (b,c).

Multiple localizations generated by repeated blinking of the same molecule were combined into a single localization by the Distance Distribution Correction (DDC) algorithm recently devised by Bohrer et al. [37]. DDC infers the true pairwise distance distribution of different fluorophores directly from the temporal sequence of images by considering distances between localizations that occur with a time lag much longer than the average fluorescence survival time. Accordingly, DDC removes blinking artifacts without additional calibrations, affording a close estimate of true spatial organization of emitters. Final, DDC-cured dSTORM maps were characterized by a density of 19±3 emitter/µm^2^ (average±SE, #29 cells), with clear heterogeneous arrangement of single emitters in nanoscale clusters (**Figure 6a,b**). The comparison between localization (pre-DDC) and emitter (post-DDC) densities revealed that a single emitter was localized 2.4 times on average. Given our exposure time of 30 ms per frame, this value corresponded to 72 ms as the average lifespan of fluorescence emission of AlexaFluor647 in typical dSTORM conditions, this value being in excellent agreement with the 70-80 ms reported by Bohrer et al. [37].

A quantitative description of the nanoscale organization of PD-L1 was provided by Ripley’s cluster analysis on cleaned dSTORM data. Ripley’s method leverages a second-moment property of the single molecule spatial map to perform a statistical test between the actual point distribution and that associated with the isotropic dispersion of molecules in space [52]. More specifically, Ripley’s method calculates a global function H(r) for increasing distances *r* from every single molecule in the *xy* image molecules, and *H(r)*<0 indicates spatial dispersion of molecules. The *r_m_* value plane: H(r)>0 indicates clusterization, H(r)=0 indicates a uniform distribution of corresponding to the maximum of H(r) is a proxy of cluster size [38; 39].

Cells were analyzed by Ripley’s method in the interval 1-1,000 nm. H(r) was always calculated in large membrane areas (100-850 μm^2^) to afford a representative result of PD-L1 membrane distribution. In all cases, H(r) grew sharply within the first 100 nm, reached a maximum, and slowly declined afterwards, suggesting the existence of nanoscale clusters (**Figure 7a**). A Monte Carlo algorithm was applied to correct the experimental Ripley’s function for the localization precision of the dataset, using the value nm. It is worth noting that the DDC procedure effectively decrease σ_xy_, due to merging of those localizations that pertain to the same molecules. Thus, albeit minor, the correction applied to the experimental Ripley’s function was interpreted as a maximum estimate.

**Fig. 7.**
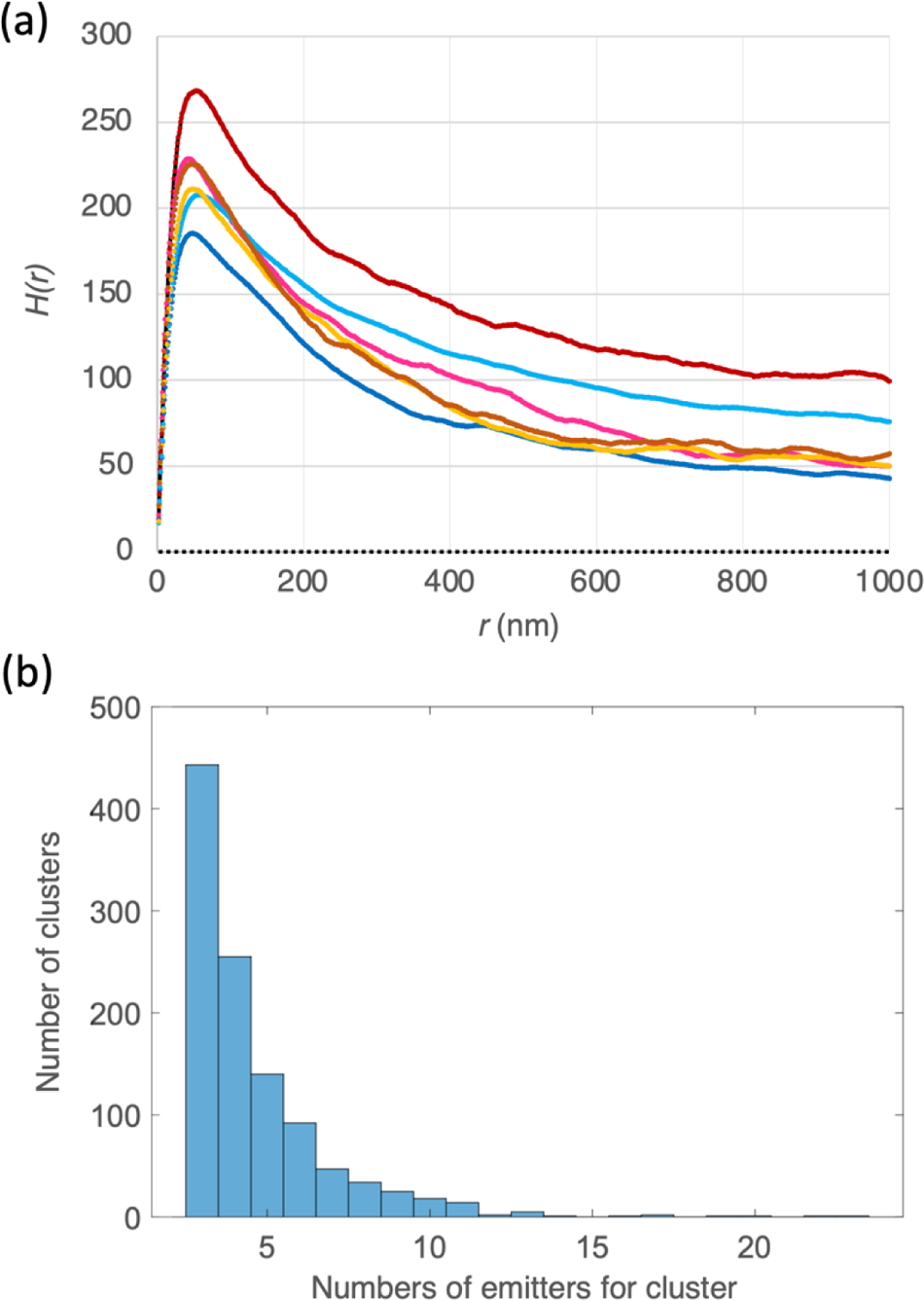
Quantitative cluster analysis of single molecule maps. (a) Plot of the Ripley’s function H(r) for 6 different cells. The dashed black line for H(r)=0 refers to isotropic distribution of molecules. (b) Histogram of cluster density of emitters after Getis and Franklin’s local point pattern analysis and morphological evaluation. Note that cluster must have ≥3 emitters to be considered as such.

We found out <r_m_> = 42.7±1.0 nm (average±SE, #29 cells). Assuming the scaling factor of 1.3 inferred by Ruan et al. [39], this value converts into a minimum cluster radius *r_c_* = 32.8+0.8 distribution with the size (*d_Ab_*) of αPD-L1-647, which we approximated to 7-10 nm by 32.8±0.8 nm. Of note, this measured radius is a convolution of the PD-L1 aggregation halving the enlargement data reported for a primary-secondary Ab system in [53]. From this value, a simple heuristic rule was applied to estimate the convolution effect on the actual radius *r^0^_c_* of the cluster [54]:

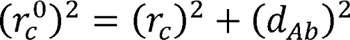

Setting *d_Ab_*= 10 nm, we obtained *r^0^_c ≌_* 30 nm, i.e. the finite size of the antibody produced only a minor increase to the real cluster radius.

We further analyzed the nanoscale aggregation of PD-L1 by creating quantitative clustering maps according to Getis and Franklin’s local point pattern analysis [41; 42]. In short, we defined a reasonable clustering threshold of H(r) for r= 40 nm (the integer closer to and we generated a binary map containing clustered (above threshold) and non-clustered (below threshold) areas for each cell dataset (Figure 6c).

Morphological analysis of the binary maps returned crucial properties such as the density of localizations in the cluster (*d*_c_) and the % of overall localizations belonging to Morphological analysis of the binary maps returned crucial properties such as the participate in nanoclusters, as revealed by <F> =65.4±1.9% (average±SE). We also found clusters (f). Significantly, more than 50% of membrane PD-L1 molecules were found to <*d_c_*> =5.2±0.1 (average±SE) localization/cluster (**Figure 7b**). Assuming a 1:1 binding stoichiometry between αPD-L1-647 and PD-L1, this value was used to estimate the number of clusterized PD-L1 molecules by considering the degree of labeling of the fluorescent antibody. A standard spectrophotometric quantification method yielded 1.08 as the degree of labeling of αPD-L1-647, positing 5 PD-L1 molecules per nanoaggregate. Notably, this value represents a minimum estimate of the molecular density, as we must consider the impact of incomplete antibody decoration of PD-L1, a well as the non-exhaustive detection of all possible emitters.

## DISCUSSION

In the last few years, PD-L1 has become one of the most important therapeutic targets in oncology [7], on account of its remarkable standing as a cellular checkpoint of immune escape in many cancers [5; 6]. The molecular details of PD-L1 binding to its immune receptor PD-1 are well described. Yet, little is known about the arrangement of PD-L1 on the plasma membrane, a feature that must play a crucial role in the engagement of immune checkpoint partner PD-1 and could represent an interesting therapeutic target. The membrane organization of PD-L1 could also help the understanding of some other intrinsic mechanisms leveraged by PD-L1 to regulate relevant aspects of tumor progression [10].

Because of the palmitoylation of Cys272, a residue located in the cytoplasmic tail of PD-L1, it has been suggested that PD-L1 might be engaged in the raft regions of the cell membrane, given the strict association between palmitoylated proteins and lipid rafts [26]. Some experiments on ex-vivo membranes seem to support this hypothesis [27]. In this work, we set out to clarify the alleged raft engagement of PD-L1 by investigating the membrane organization of PD-L1 following a multiscale imaging approach, which leverages both diffraction-limited and super-resolution fluorescence microscopy techniques.

NSCLC constitutes one of the most intense clinical research areas on PD-L1, owing to the success of first-line immunotherapy in a significant subset of cases [8], albeit most patients eventually develop resistance. NSCLC cell lines bearing mutations on the EGFR gene have been reported to express medium-to-high levels of membrane PD-L1, offering advantageous models to study PD-L1 in the cellular setting with high sensitivity. Accordingly, we initially screened five popular NSCLC cell lines with different mutations on EGFR by cytofluorometry. Our screening identified HCC827, characterized by the 746-750 deletion on EGFR, as the cell line with the highest expression of PD-L1.

Confocal, TIRF and ISM^++^ imaging of immunostained cells clearly demonstrated a heterogeneous organization of membrane PD-L1 into submicron aggregates (Figures 2-3). Additionally, PD-L1 was found to colocalize strongly with two classical hallmarks of lipid raft regions, glycosylphosphatidylinositol (GPI) and caveolin-1 (Cav-1), advocating for the engagement of PD-L1 in membrane rafts. In keeping with this hypothesis, membrane cholesterol depletion by MBCD led to significant loss of membrane PD-L1, albeit the heterogeneous organization morphology onto the PM was maintained.

Lipid rafts are dynamic biological platforms occurring at nanoscale [23]. Diffraction-limited confocal and TIRF microscopies are unable to reach spatial resolutions below 200 nm [29]. This limit is only partially alleviated by ISM^++^, whose resolution does not exceed the 100-140 nm mesoscale [34]. Thus, we applied single molecule localization microscopy (dSTORM approach) in the TIRF mode to directly observe the alleged nanoscale organization of PD-L1. For this analysis, we took advantage of an anti-PD-L1 IgG conjugated, on average, with one Alexa647 fluorophore. Under the reasonable assumption of 1:1 stoichiometry between PD-L1 and the IgG, localization maps of single Alexa647 emitters were interpreted as spatial maps of single PD-L1 molecules. Ripley and G&F cluster analysis of the localization maps revealed that PD-L1 assembles in nanoscale aggregates of about 30-40 nm radius, each containing at least 5 proteins on average. Of note, G&F analysis also revealed that more than half (∼65%) of the membrane PD-L1 pool aggregates in nanoclusters, in keeping with the incomplete colocalization of the protein with raft markers. These findings strongly support a model of PM organization where a large pool of PD-L1 is sequestered in nanosized raft regions.

## CONCLUSIONS

In conclusion, we characterized the relationships of PD-L1 with lipid rafts in the plasma membrane by taking advantage of a toolbox of fluorescence microscopy techniques operating at different spatial resolutions. As a cellular model, we selected the NSCLC EGFR-mutated cell line, HCC827, characterized by high membrane expression of PD-L1. For the first time, the actual engagement of a large (65%) PD-L1 pool in the raft regions was directly demonstrated in the unperturbed PM. This picture was supported by experimental results: PD-L1 largely colocalizes with raft membrane markers and it aggregates as nanoclusters of 30-40 nm radius, in excellent agreement with the reported size of PM raft regions. Of note, our membrane-specific fluorescent labeling of PD-L1 proved effective in all imaging conditions.

Raft engagement could disclose unreported biological functions of PD-L1. Indeed, following the preliminary results of Zhang [27], it is tempting to speculate that raft engagement may be crucial for the immune checkpoint connected to the PD-L1/PD-1 interaction. This knowledge could help the development of new immuno-oncology strategies. Additionally, the non-clustered pool could be related to other non-immune functions of the protein, most of which were recently discussed by Deng et al. [10] Further experiments are under way to clarify this point.

## Acknowledgments

Dr. Michele Oneto (IIT Nanophysics) are gratefully acknowledged for technical assistance and support.

## DECLARATIONS

### Author Contributions

Barbara Storti and Ranieri Bizzarri conceived and designed the study. Material preparation, data collection and analysis were performed by all authors. The first draft of the manuscript was written by Martina Ruglioni, Simone Civita, Barbara Storti and Ranieri Bizzarri and all authors commented on previous versions of the manuscript. All authors read and approved the final manuscript.

### Data avaliability

data will be made available upon reasonable request.

## Footnotes

1 https://tools.thermofisher.com/content/sfs/manuals/mp20173.pdf.

2 https://sites.broadinstitute.org/ccle/

